# Robust Gap Repair in the Contractile Ring Ensures Timely Completion of Cytokinesis

**DOI:** 10.1101/052704

**Authors:** AM Silva, D Osório, AJ Pereira, H Maiato, IM Pinto, B Rubinstein, R Gassmann, IA Telley, AX Carvalho

## Abstract

Cytokinesis in animal cells requires the constriction of an actomyosin contractile ring, whose architecture and mechanism remain poorly understood. We use laser microsurgery to explore the biophysical properties of constricting contractile rings in *C. elegans* embryos. Laser cutting causes rings to snap open, which is a sign of tension release. However, instead of disintegrating, ring topology recovers and constriction proceeds. In response to severing, a finite gap forms that is proportional to ring perimeters before cutting, demonstrating that tension along the ring decreases throughout constriction. Severed rings repair their gaps by recruiting new material and subsequently increase constriction rate and complete cytokinesis with the same timing as uncut rings. Rings repair successive cuts and exhibit substantial constriction when gap repair is prevented. Our analysis suggests that cytokinesis is accomplished by contractile modules that assemble and contract autonomously, enabling local repair of the actomyosin network throughout constriction. Consequently, cytokinesis is a highly robust process impervious to discontinuities in contractile ring structure.

## Introduction

Cytokinesis, the process that completes cell division by partitioning the contents of the mother cell to the two daughter cells, requires the assembly and constriction of a contractile ring (Green et al., 2011). The contractile ring is composed of a network of actin filaments and non-muscle myosin-II motor proteins, whose architecture and mechanism of constriction remain poorly understood. The contractile ring could be composed of a series of contractile units, (Carvalho et al., 2009; Bement and Capco, 1991) although a modular organization has so far not been observed by electron microscopy (Kamasaki et al., 2007). Alternatively, the ring could be composed of a single actomyosin bundle, which continuously self-remodels, as proposed based on work in fission yeast protoplasts (Stachowiak et al., 2014). Laser microsurgery has provided powerful insight into cellular actomyosin structures and has been used to study wound healing (Mandato and Bement, 2001), stress fibers (Colombelli et al., 2009; Kumar et al., 2006), cortex dynamics (Tinevez et al., 2009), cohesion of the epithelial tissue during cell division (Guillot and Lecuit, 2013; Founounou et al., 2013; Herszterg et al., 2013) and shape changes during tissue morphogenesis (Munjal et al., 2015; Collinet et al., 2015). Here, we combine laser microsurgery with live-imaging in the *C. elegans* early embryo to probe the biophysical properties of constricting contractile rings during cytokinesis.

## Results and Discussion

In the 4-cell embryo, 2 cells (ABa and ABp) undergo contractile ring constriction parallel to the focal plane, which provides an end-on view of the constricting ring and facilitates the monitoring of shape change and ring component dynamics (Fig. S1A). Furthermore, these cells are relatively large with a contractile ring perimeter of 55.8 ± 0.4 μm (mean ± 95% confidence interval) at constriction onset and divide with highly reproducible kinetics, enabling spatiotemporal measurements and analysis (Fig. S1C, D). The rate of ring constriction is roughly constant until the ring approaches the spindle midzone, after which ring closure slows down until constriction completes (Fig. S1D) (Carvalho et al., 2009).

Using nanosecond laser pulses, we severed rings of ABa and ABp cells expressing myosin^NMY–2^::GFP, a fluorescent non-muscle myosin II heavy chain that localizes in the ring and in the remaining cell cortex, at different stages of constriction (Fig. S1A, B). Contractile rings exhibited a fast mechanical change of strain, snapping open immediately after the cut. Surprisingly, rings neither regressed completely nor showed signs of disintegration. Instead, after finite retraction of the severed ends, the gap was rapidly filled in with myosin^NMY–2^::GFP and ring constriction resumed (Fig. 1A, B). Severed rings always successfully completed constriction (n=75) and embryos continued to develop (7 out of 11 tracked embryos hatched and the remaining embryos arrested during elongation, a late stage of embryogenesis). A striking finding is that despite locally destroyed ring topology, the resulting ring fragment retained most of its shape, which implies that it is anchored to the plasma membrane. Moreover, a mechanism is in place to repair gaps at any stage of constriction, invariably allowing for successful conclusion of cytokinesis.

**Figure 1.**
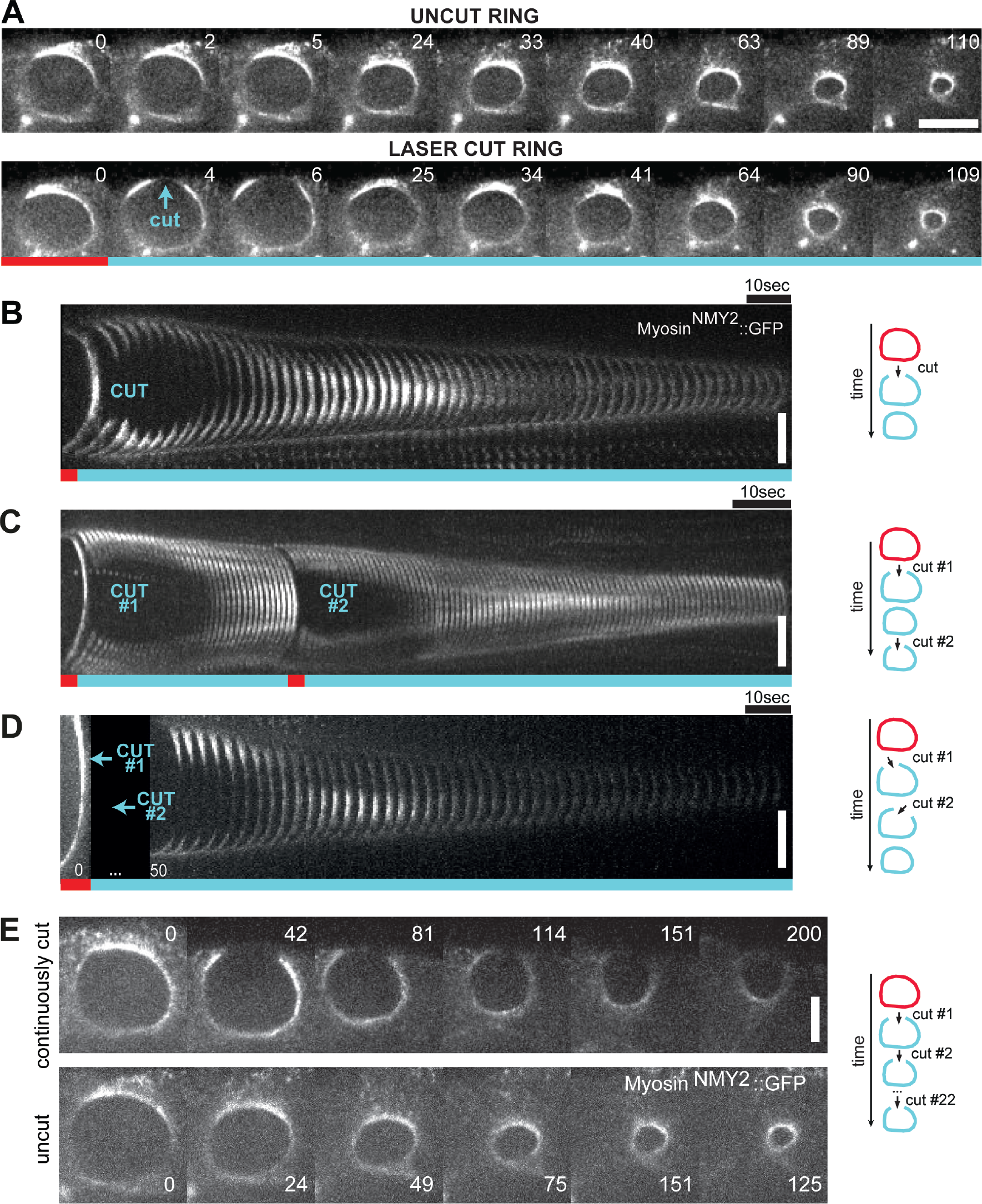
Behavior of contractile rings severed by laser microsurgery during constriction. **A.** End-on view stills from time-lapse imaging series of constricting contractile rings in ABp cells expressing myosin^NMY–2^::GFP. The contractile ring snaps open after the laser cut (cyan arrow, frame 4), the gap is repaired (frame 25), and ring constriction resumes to completion. Time is in seconds; time zero denotes frame before the laser cut. **B-D.** Kymographs of ring regions cut by laser microsurgery built as depicted in Fig. S1E. Time interval between frames is 2.32 seconds in B, 1 second in C and 3.2 seconds in D. Rings were subjected to a single cut (B), or two sequential cuts with second cut either hitting the same region (C) or a neighboring region (D). Time-lapse acquisition in D resumed after both cuts were executed (missing frames are indicated by a black box). **F.** Stills of a constricting ring with a persistent gap, which was maintained by iteratively cutting the same site. Time is in seconds, time zero denotes frame before the laser cut. Scale bars, 5 μm.

To assess the robustness of contractile ring repair, we performed sequential cuts on the same constricting ring. In one set of experiments (n=6, Fig. 1C), the second cut was made at the site of the first cut after the gap had been repaired. In another set of experiments, sequential cuts were made in spatially separate regions 20-25 s apart (n=8, Fig. 1D). Similar to rings that were cut once, rings cut twice with either cutting regime repaired the gaps and completed constriction. These results demonstrate that ring gap repair is history-independent and a locally autonomous process that is not influenced by events in adjacent regions of the ring.

To test whether gaps in contractile ring structure must be repaired in order for constriction to proceed, we made 22 consecutive cuts on the same side of a constricting ring, thus preventing it from resealing for 300 s. Strikingly, the ring fragment shortened from 30 μm to 10 μm during this time (Fig. 1E). Although the ring fragment constricted at a reduced rate compared with uncut control rings, this experiment shows that a continuous contractile ring structure is not a prerequisite for constriction.

Next, we followed the initial response of severed rings with high temporal resolution by resuming time-lapse imaging 2.6 s after the laser cut and then acquired one frame per second (or 7 z-planes per 2.32 s). We analyzed constricting rings with perimeters ranging from 47.2 to 11.2 μm immediately before the cut, hereafter referred to as initial perimeter. In these experiments, we were able to accurately follow the severed ends, allowing us to determine the gap size and the length of the cut ring (arc length) for 10 s after the cut (Fig. 2A–E). The response to the laser cut resembled a viscoelastic mechanical relaxation, similar to what has been previously described for rings in an epithelial tissue (Guillot and Lecuit, 2013). At least two phases of gap formation after microsurgery were detectable within our temporal resolution: A fast phase of elastic recoil, and a slowdown (“creep”) phase that lasted for ~6 s after the cut, when the gap reached a maximum size of 20.2 ± 0.9% of the initial perimeter (n=24, Fig. 2C, S2A). A fraction of the fast gap formation (<1 μm) is due to material removal caused by ablation with a diffraction limited laser spot. After the gap widening phase, the gap size remained roughly constant for ~4 s (Fig. 2C, S2A) and subsequently decreased in a phase of repair. Throughout this multiphasic response the arc length of the open ring continued to decrease at a rate comparable to uncut rings (0.30 μm/s for group I versus 0.27 μm/s for uncut rings, Fig. 2D). We note a tendency for initially larger rings to shorten more during the elastic response. All this indicates that the presence of the gap did not impact the ability of the open ring to constrict at normal speed. It also suggests that either some other mechanical component like the cell cortex or the plasma membrane is assisting ring constriction, or that the remaining ring arc is still able to do work, which would imply that it is under tension.

**Figure 2.**
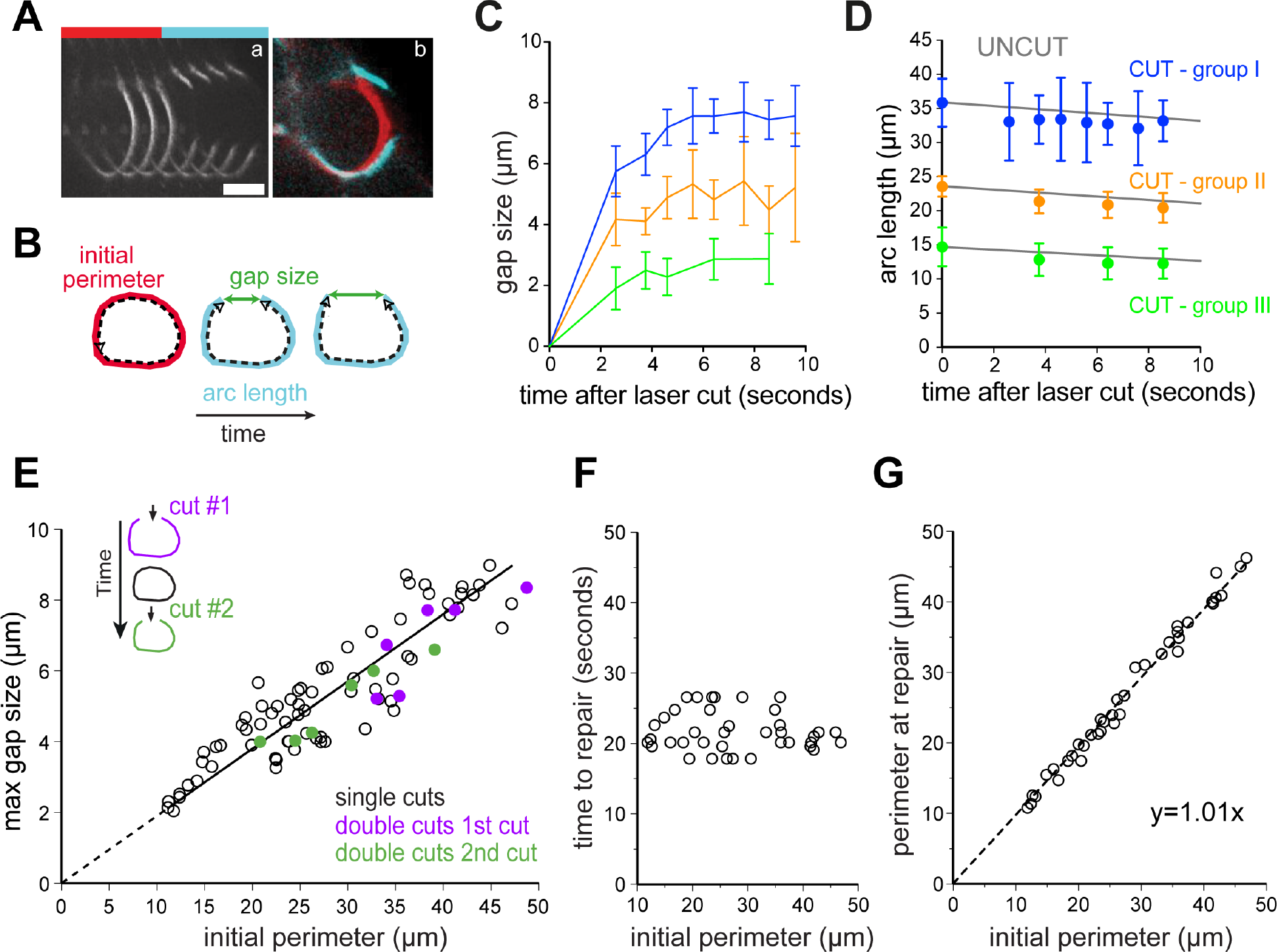
Analysis of gap formation and repair after laser microsurgery in contractile rings. **A.** Kymograph of cut ring region showing the snapping event and straightening of the severed ends (a). Time interval between frame 3 and 4 is 2.6 s; other frames are 1 second apart. Overlay (b) shows frames immediately before (red) and after (cyan) the cut. **B.** Schematic defining initial ring perimeter before the cut, arc length, and gap size. **C, D.** Gap size (C) and arc length (D) measured during the first 10 s following the cut (n=24). Cut rings were divided in three groups with initial perimeters averages of 35.8 μm (group I), 23.6 μm (group II), and 14.6 μm (group III). Grey lines in D represent the average perimeter decrease for uncut rings (data as in Fig. S1C). Error bars represent the 95% CI of the mean. **E.** Maximal gap size plotted against initial perimeter for rings cut once (black circles, n=69) or twice as in Fig. 1C (first cut in purple, second cut in green; n=6). Maximum gap size and initial perimeter are proportional (R^2^=0.81; r=0.90; p<0.0001). F. Time between laser cut and completion of gap repair plotted against initial perimeter (n=38). G. The perimeter at the time of completion of repair plotted against initial perimeter, showing a strong correlation (n=38; R^2^=0.99; r=0.99; p<0.0001).

Immediately after the laser cut, the severed ends no longer followed the curvature of the ring, instead adopting a straight conformation (Fig. 2A). Instantaneous and local relaxation of cortical tension is likely responsible for pulling the severed ends outward, as it has been proposed that contractile ring constriction is influenced by the tension in the remaining cell cortex (Zhang and Robinson, 2005; Turlier et al., 2014; Sain et al., 2015). Ring snapping and straightening of severed ends were highly consistent responses to laser microsurgery (71 out of 75 cut rings) and was also observed in rings marked with other contractile ring components including lifeAct::GFP (marking actin) and GFP::septin^UNC–59^ (Fig. S2B). Moreover, we observed a corresponding deformation of the plasma membrane at the cut site using GFP::PH^PLC1δ1^ as a marker (Fig. S2C)(Audhya, 2005), which revealed local relaxation of the plasma membrane. The curvature of the plasma membrane recovered 10-15 s after the cut (Fig. S2C).

A mechanical model for cytokinesis in the early *C. elegans* embryo postulated that line tension in the ring is proportional to ring radius during constriction in the quasi-static approximation (Sain et al., 2015). In our laser microsurgery experiments, gap size is primarily a readout of local tension within the contractile ring and therefore we expected to find a linear relation between gap and ring sizes. Strikingly, we found a significant proportionality between maximum gap size and initial perimeter (R^2^=0.80, n=69; Fig. 2E). The observation that gap size scales with initial ring perimeter matches qualitatively the prediction of the static model and demonstrates that ring tension decreases throughout ring constriction. In cases where constricting rings were cut sequentially (Fig. 1C), the gap size after the second cut was again proportional to the initial perimeter before the second cut. This demonstrates that closing the gap after the first cut had reestablished line tension locally within the ring (Fig. 2E).

Following gap widening (i.e. 6 s after the cut), a 12-second period followed when we could not accurately identify the severed ends. During this time, gaps were completely filled in with myosin^NMY–2^::GFP, lifeAct::GFP and GFP::septin^UNC–59^ (Fig. 1A, B, S2B) such that 21.9 ± 1.0 s after the laser cut all gaps were undetectable and a closed ring topology was re-established (Fig. 2F). We considered this to be the time point when gap repair was complete. Therefore, the time required for cut rings to completely repair gaps ranging from 2.1 to 8.9 μm was constant (Fig. 2E, F). Interestingly, the perimeter at which repair completed matched the initial perimeter (Fig. 2G), indicating that the previous line tension was established. The invariable time to complete ring repair suggested that contractile ring components were recruited along their entire length of the gap, as gap closure from severed ends would be predicted to take longer for larger gaps. In agreement with this, myosin intensity profiles along the circumference of the ring revealed no significant lateral mobility from within the ring towards the gap (Fig. S2F). Analysis of a minority of cut rings (2 out of 71), in which the first attempt at gap repair failed and the plasma membrane regressed more than usual, lent further support to this idea. In these two cases, myosin recruitment could be observed throughout the membrane at distinct foci (arrows in Fig. S2D), and the gap was completely filled in with myosin^NMY–2^::GFP 20 s after the first failed repair (35 s after the cut). We conclude that within 22 s of being severed by laser microsurgery, constricting contractile rings have repaired their gaps from multiple sites along the exposed plasma membrane.

To examine the impact of the repair process on the kinetics of ring constriction, we analyzed the rate of change in ring perimeter after repair. Remarkably, this analysis revealed that cut rings took the same time to constrict to completion as uncut control rings (Fig. 3A). Inspection of individual traces revealed that the constriction rate of cut rings was on average 35% higher than that of uncut control rings (n=15; Fig. 3B). Cut rings eventually caught up with uncut control rings likely because of the proximity to the spindle midzone. The spindle midzone has been shown to slow down contractile ring constriction (Carvalho et al., 2009), suggesting that it imposes a limit on the interval during which repaired rings can close with an increased constriction rate. In agreement with this, repaired rings with larger initial perimeters, which started further away from the spindle midzone, closed with an increased constriction rate for a longer time before their perimeter caught up with the average perimeter of uncut control rings (R^2^=0.59; Fig. 3C). Along with accelerated constriction we observed transient myosin accumulation in the repaired region above the average myosin levels in that region before the cut. Myosin levels peaked 45.2 ± 1.9 s post-cut and then decreased and stabilized at ~0.6 times the initial value, 73.9 ± 4.7 s post-cut (Fig. 3D, S3B, F). Actin also accumulated in the same region with levels peaking 26.2 ± 1.5 after the cut (Fig. S2B, S3B). Interestingly, the increase in myosin levels beyond the stabilization level roughly coincided with the time of repair completion, always occurring within the interval of increased constriction rate (Fig. 3B, D and S3F). We conclude that, following complete gap repair and recovery of ring topology, cut rings transiently recruit an excess of myosin to the repair site and speed up constriction, allowing them to complete cytokinesis in the same time as uncut rings.

**Figure 3.**
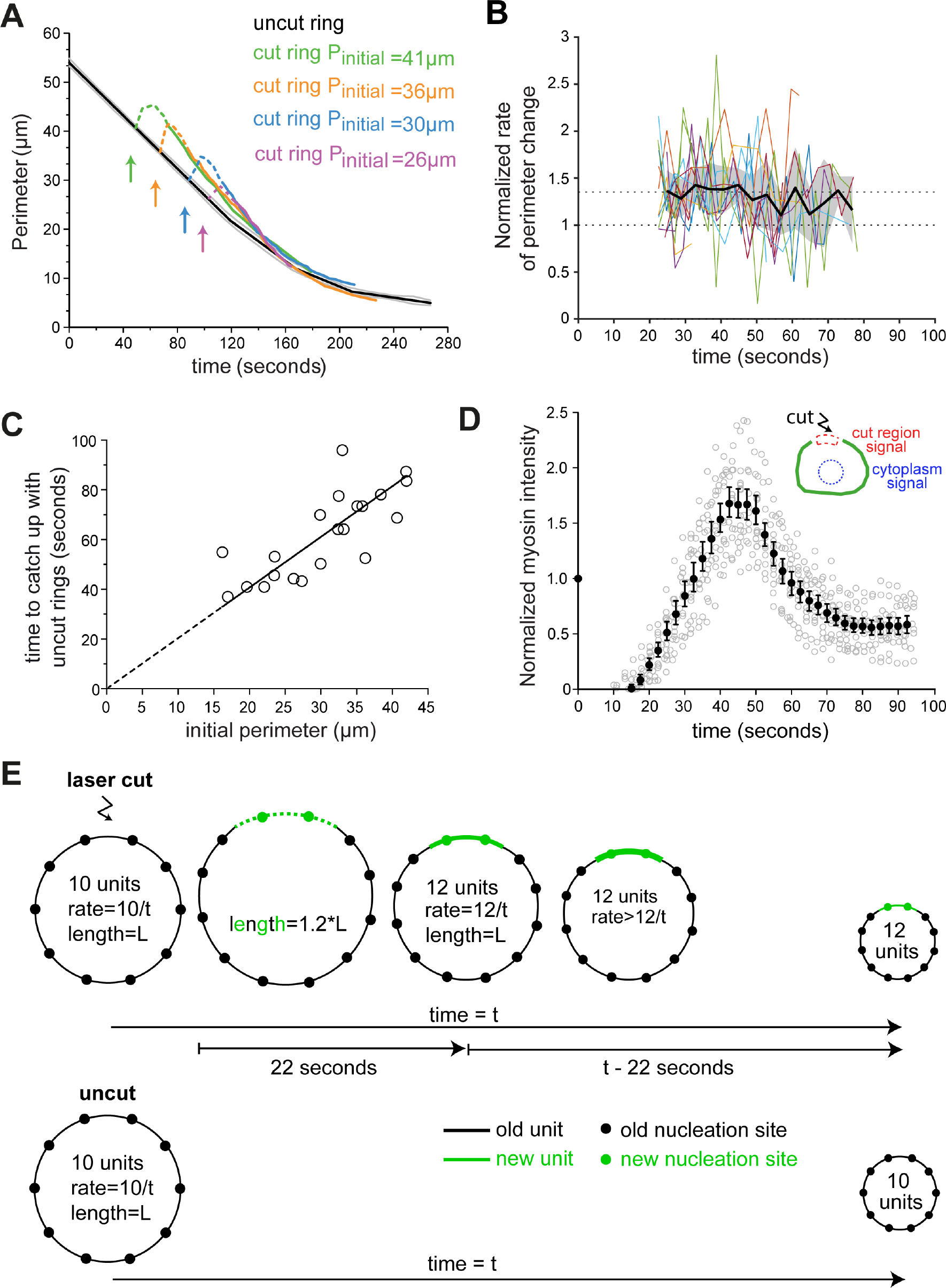
Repaired rings close faster and catch up with uncut control rings. **A.** Perimeter versus time plot of four cut rings with initial perimeters (P_initial_) of 41 μm, 36 μm, 30 μm and 26 μm. Dashed portion of colored lines corresponds to time points from 0 to ~22 s, for which part of the measured perimeter corresponds to the outline of cortical myosin^NMY–2^::GFP labelling. Average perimeter for uncut rings is in black with grey lines indicating the 95% CI (data as in Fig. S1C). Arrows indicate moment of laser cut. **B.** Normalized ratio of perimeter change over time after cut (n=15; see Fig. S1F and methods for details). Colored lines represent individual examples and the black solid line shows the average. Dashed lines indicate normalized ratios of 1 and 1.35 (average of normalized rates above 1 over the time period of 22-55 s after the laser cut). Grey area represents the 95% CI of the average curve. **C.** Time required for cut rings to catch up with the perimeter of uncut rings plotted against initial perimeter (n=21; R^2^=0.59; r=0.76; p<0.0001). **D.** Normalized myosin fluorescence intensity levels plotted against time after cut for rings with initial perimeters between 16 and 42 μm (see Fig. S3A for method). Black dots represent average intensities (see methods for details) with 95% CI error bars; grey circles represent individual measurements. **E.** An increase in constriction rate proportional to the gap size can be explained by the contractile unit model. When a ring is cut, the plasma membrane retracts at the site of the cut and contractile units (green) assemble *de novo* at nucleation sites (green circles) distributed along the plasma membrane. This increases the total number of units in the ring and therefore the overall rate of constriction after gap repair relative to uncut control rings. We propose that subsequent myosin hyper-accumulation (thick green line) transiently increases tension locally along the new units and promotes faster shortening of the repaired ring, leading to an overall higher rate of perimeter constriction than expected by serial incorporation of new units.

Our analysis of contractile ring dynamics after laser microsurgery provides strong support for the previously proposed model that constriction is powered by a series of contractile units of equal size that locally constrict the plasma membrane at a constant rate and make additive contributions to the overall constriction rate (Carvalho et al. 2009). We propose that contractile units are connected to one another such that the contractile ring is able to sustain tension. The functional autonomy of individual contractile units explains the observation that constriction proceeds despite the presence of a gap in the ring circumference. Furthermore, units must be anchored to the membrane and/or cortex and sustain tension in the severed ring, which explains why a severed ring does not fall apart and exhibits only minimal change in curvature. Our measurements of gap size after laser cutting reveal that tension within the contractile ring decreases during constriction, which is in agreement with a previously proposed mechanical model for cytokinesis in *C. elegans* embryos (Sain et al., 2015). This will constitute an important parameter to feed into future mathematical and physical models of cytokinesis. We show that at any time during constriction, gaps varying in size up to 4-fold are efficiently repaired with identical kinetics, and that multiple sites of myosin recruitment are present during the repair of larger gaps. We speculate that those discrete recruitment sites correspond to nucleation foci on the ‘naked’ plasma membrane exposed after a laser cut, from which new contractile units are assembled. Such nucleation sites may correspond to points where contractile units are anchored to the plasma membrane, allowing for line tension transmission between neighboring units and lateral tension transmission to the membrane and remaining cortex.

The rapid kinetics of gap repair highlights the remarkable capacity for local remodeling of contractile ring structure throughout constriction, which makes cytokinesis a highly robust process unlikely to fail once the contractile ring has assembled. Importantly, the contractile units assembled *de novo* at the site of the gap increase the total number of units in the repaired ring. As the overall rate of ring constriction is proportional to the total number of units, the incorporation of additional units into repaired rings provides an explanation for the proportional increase in constriction rate (Fig. 3E). In addition, it is conceivable that local hyper-accumulation of myosin reflects parallel incorporation, increasing the cross-sectional force capacity in the newly assembled units. Myosin hyper-accumulation would increase ring tension transiently and thereby work against resistive mechanical components, such as viscous drag and surface tension, in a dynamic force balance at a faster shortening rate. Thus, the acceleration of repaired rings may be promoted by both gain of additional contractile units and transient increase of local myosin density at the site of the laser cut. However, we note that in some cases myosin levels drop to stabilization levels before ring slowdown (Fig. S3D–F). This argues against higher myosin density and higher force capacity to be the sole cause for increased constriction rate.

The reason why myosin hyper-accumulates in the repair region remains unclear. Interestingly, a transient local overshoot of myosin was also reported after bleb retraction in metaphase cells (Biro et al., 2013). We hypothesize that, in analogy to what occurs after bleb retraction, the local spike of myosin in the contractile ring is triggered by local deformation of the plasma membrane at the cut site. Finally, previous experiments in *Xenopus* oocytes showed that contractile rings forming around plasma membrane wounds can also snap open and suffer limited recoil (Mandato and Bement, 2001). It would be interesting to assess whether rapid gap repair, local actomyosin hyper-accumulation and ring acceleration are also observed in this context, as these responses may be universal features of actomyosin contractile rings.

## Materials and Methods

### *C. elegans* Strains

Strains used in this study are listed in Table S1 and were maintained at 20 °C.

### Live imaging

Live imaging was performed on 4-cell stage embryos mounted on microfabricated chambers (Carvalho et al., 2011) at 23 °C.

### Laser microsurgery

Laser microsurgery in constricting contractile rings was performed by using the imaging objective (1.4 NA) to focus the second-harmonic beam (532 nm wavelength) of a pulsed Nd:YAG laser (Elforlight, model FQ–500–532) which is injected into the microscope stand through a secondary filter turret (‘stage-up kit’). Pulse width is 10ns and pulse energy is 1.5–2 J with the sample being exposed to a 1 kHz pulse train for 800 ms. The laser spot at the sample is a PSF-shaped ellipsoid ~0.5μm wide in the x-y plane and ~1.2 μm along the optical axis. The scale of the ablation volume is comparable but larger than the laser spot volume, which is due to the requirement that the laser intensity surpasses the ablation threshold intensity across the full contractile ring width. A spinning disk confocal system with a CSU-X1 confocal spinning disk head (Yokogawa) was mounted on an inverted microscope (TE2000U; Nikon) equipped with an electron multiplying charge coupled device camera (iXon 897+, Andor Technology), a 100x 1.4 NA oil-immersion plan-apochromatic objective (Nikon) and a 488 nm laser and a 561 nm laser. Laser shuttering and microscope hardware were controlled using NIS Elements software (Nikon) and a DAC card (National Instruments, PCI 8733). Laser microsurgery was performed at different stages of ring constriction at ring perimeter between 11 μm and 49 μm. Image acquisition was initiated before the ring reached the desired stage of constriction. By that time the embryo was moved in order to have the ring positioned in the laser microsurgery target site and the laser cut was performed. It took on average 2.6 s for the laser cut procedure, during which image acquisition was paused. For the experiments in figure 1C–E, two or more sequential cuts were made in the same constricting ring, either in the same region with the second cut being inflicted where the repair from the first cut occurred (Fig. 1C), or in neighboring regions along the ring (Fig. 1D). In order to obtain cuts in neighboring regions, the cuts were inflicted approximately 20 s apart. In this case, image acquisition was stopped before the first cut and only resumed after the second cut. In order to keep the ring continuously open, 22 consecutive cuts were performed in the same region of the ring (Fig. 1E).

To monitor contractile ring behavior immediately after laser microsurgery, a single image was acquired in the 488 nm channel every second; to follow the repair and constriction after laser microsurgery, 5.0×0.5 μm, 7.0×0.5 μm or 7.0×1.0 μm z-stacks were collected in the 488 nm channel every 1.7 s, 2.32 s and 3.2 s respectively (fastest acquisition mode). To monitor contractile ring constriction of uncut rings, 7.0×0.5 μm or 11.0×1.0 μm z-stacks were collected every 2.32 s or 10 s, respectively.

Plasma membrane behavior after laser cutting was monitored in 4-cell embryos expressing a plasma membrane probe (pleckstrin homology domain PLC1 delta1) fused with GFP and myosin^NMY-2^::mCherry. The mCherry signal enabled ring visualization until the laser cut was performed. After the cut, the mCherry signal was photobleached and the behavior of the membrane probe in the 488 nm channel was followed by collecting a 5.0×1.0 μm z-stack every 4.3 s (Fig. S2C).

## Image analysis, quantifications and statistics

All measurements and image processing were done using Fiji (Schindelin et al., 2012; Schneider et al., 2012). Z-stacks were projected using the “maximum intensity projection” tool (Fig. 1, 2A, S1A, S2C, D). Images within each panel were scaled equally. Kymographs shown in figures 1B–D and 2A were obtained using the 3D project tool. A region in maximum intensity projections or single sections that included the area of the laser cut was selected to create the kymographs (Fig. S1E). Linear regressions and statistical analyses were performed with GraphPad Prism 6.0 (GraphPad Software, Inc., San Diego, CA). R^2^ denotes the coefficient of determination of the linear regression; r denotes Pearson’s correlation coefficient. All error bars represent the 95% CI of the mean. Rate calculations were done using Matlab (MathWorks Inc., Natick, MA).

<u>1. Measurement of contractile ring perimeter and mean constriction rate during ring constriction</u>

All measurements were performed on constricting rings of ABa or ABp cells expressing myosin^NMY–2^::GFP, with the exception of some of the measurements presented in Fig. S3B, which were done in cells expressing lifeact::GFP. For uncut rings, the ABa and ABp ring perimeter was determined by manually tracing the ring outline in maximum intensity projections of z-stacks using the Segmented Line tool in Fiji and reading the length of the total segmented line. Data from multiple rings were temporally aligned and averaged by calculating the arithmetic mean of ring perimeters within a time frame (Fig. S1C). We could resolve four phases of the time-perimeter curve by fitting each phase to a linear trend and obtaining correlation values R^2^ close to 1.00. The equations of the linear trends were: y = −0.27x + 46.9 for ring perimeters between 47.1 μm and 22.8 μm, y = −0.20x + 40.7 for ring perimeters between 20.6 μm and 12.5 μm, y = −0.11x + 28.0 for ring perimeters between 11.1 μm and 7.7 μm, and y = −0.04x + 14.6 for ring perimeters between 7.0 μm and 5.0 μm (Fig. 2D, 3A, S3F). These functions were used to align the perimeter curves from cut rings to those of uncut rings.

The rate of constriction of individual uncut rings was calculated for pairs of consecutive time points by dividing the difference in perimeter by the time interval (first order difference method) (Fig. S1D). Data from multiple rings were averaged (Fig. S1D). The mean of data points that fell in overlapping 5-μm intervals was calculated and plotted against the perimeter at the center of each interval.

For cut rings, ring perimeter was determined as described above for uncut rings. In the period of time from the cut until repair completion (0–22 s), a portion of the measured perimeter corresponded to the outline of the cortex as judged by cortical myosin^NMY-2^::GFP labelling (dashed line in Fig. 3A). Individual cut ring perimeter traces were aligned with the fit linear trends for uncut rings relative to the time point just before the cut (Fig. 2D, 3A, S3F).

<u>2. Measurement of gap size, arc length of the severed ring, and time of gap repair</u>

Gap size measurements were performed in single sections or maximum intensity z-projections by manually tracing a straight line between the severed ends for each time point during the ~10 s after laser microsurgery (Fig. 2B, C, S2A). After the first 10 s, the gap started being filled with contractile ring material and measuring its size became impossible. Gap size was normalized to the ring perimeter before cut (initial perimeter) in Figure S2A. The arc length of the open ring within the first 10 s after the cut was measured by manually tracing a line from the left severed end to the right severed end along the ring (Fig. 2B, D). The maximum gap size was plotted against the initial perimeter in Figure 2E. The time of gap repair was defined as the interval between the time point just before the laser cut and the time point when the gap was completely filled in with myosin^NMY–2^::GFP (Fig. 2G).

<u>3. Measurement of instantaneous ring constriction rates after laser cut</u>

The instantaneous rate of change of perimeter Δ_t_p was determined with the first order difference method for time points after completion of repair and perimeters above 19 μm (at this point rings approach the spindle midzone and slowdown; Carvalho et al., 2009). The difference in perimeter between time points t[*i*+1] and t[*i*] was divided by the interval between those time points, for all frames *i* of a movie: Δ_t_p(t[*i*]) = (p(t[*i*+1])−p(t[*i*])) / (t[*i*+1]−t[*i*]). The rate of change of perimeter for the reference uncut ring Δ_t_P(t), and for every cut ring, Δ_t_p{*j*}(t), *j*=1…n, were calculated. P(t) represented the average of ABp rings. For cut rings, Δ_t_p{*j*}(t) was normalized to Δ_t_P where P(T) = p{*j*}(t), i.e. the time point T for which the perimeter of the uncut matched the cut ring was determined by interpolation, and the rate at this time point T was used for normalization: Ω{*j*}(t) = Δ_t_p{*j*}(t) / Δ_t_P(T), for all traces *j*=1…n (Fig. S1F). The rate Δ_t_P(T) was determined by interpolation of T within the discrete values of Δ_t_P(t[*i*]). An average curve for Ω{*j*}(t) was calculated by temporally aligning the Ω{*j*} with respect to the time point for severing, overlaying all Ω{*j*} (Fig. 3B), dividing the time axis in equal bins with size Δt = 4 s, and filling each bin with the normalized rates Ω{*j*}(*k*·Δt<t<(*k*+1)·Δt), *k*=1…M and M the number of bins. Then, the arithmetic mean of the values in each bin was calculated. Similarly, the standard error of the mean s.e.m. = s.d. / sqrt(N[*k*]), N[*k*] = number of values in bin k, was determined for all M bins, and the 95% CI was estimated as 1.96 times s.e.m. The resulting data was plotted with respect to the center of the bin, Δt/2 + *k*·Δt, *k* = 1…M.

The time interval for cut rings to catch up with uncut rings was defined as the interval between the time point just before the laser cut and the time point when the repaired ring reached a perimeter that was within the region of the uncut ring perimeter mean plus 95% CI (Fig. 3C and S3D).

<u>4. Quantification of myosin fluorescence levels after laser cut</u>

The myosin^NMY–2^::GFP average intensity level in the cut region was quantified within a box 0.7 μm thick and 2.0 μm wide drawn over the region where the cut was inflicted; the dimensions of the box were kept constant for all time points measured before and after the cut. A circular region in the cytoplasm, which decreased in size as the ring constricted, was used to quantify the average intensity of myosin^NMY–2^::GFP cytoplasmic signal. The cytoplasmic signal was subtracted from the mean intensity in the cut region and values were normalized to the average myosin intensity before the cut (Fig. 3D, S3A, F). Individual traces revealed that myosin levels increased to a maximum level and then decreased and stabilized until the end of ring constriction (Fig. 3D). The myosin^NMY–2^::GFP level profile revealed reproducible kinetics regardless of ring perimeter before the cut. Therefore, normalized profiles from multiple cut rings were aligned to the time point before cut and plotted against time after cut (Fig. 3D). The mean of data points that fell in overlapping 5 s intervals was calculated and plotted against the time at the center of each interval (Fig. 3D). The time between the laser cut and maximum myosin accumulation is shown in Fig. S3B. Quantification of actin levels in embryos expressing lifeAct::GFP was performed as described for myosin^NMY–2^::GFP (Fig. S3A). The time interval from the laser cut until the stabilization of myosin levels is plotted against the time it took for cut rings to catch up with uncut rings in Fig. S3D.

To assess the correlation between increased myosin levels in the region of gap repair and the increase of ring constriction rate, we determined the interval during which myosin levels in the repaired region were above the myosin stabilization level (Fig. S3C, D, F). The latter, which is represented by the vertical orange dashed lines in Figure S3C, was plotted against the time interval from completion of repair to catch up with uncut rings (Fig. S3E). In order to assess the extent of overlap between increased constriction rate and myosin hyper-accumulation, graphs of perimeter versus time and normalized myosin intensity versus time of individual examples were overlaid and two representative examples are shown in Fig. S3F.

## Myosin distribution around ring circumference

Myosin distribution along the ring in Fig. S2F was analyzed using Mathematica (Wolfram Research, Champaign, IL) and Image J (NIH, Bethesda, MA) with custom made plug-ins. The z-stack sum projection was computed and background was subtracted for each time frame t_i_. A 3-pixel wide polyline was manually traced along the ring for each time point and the intensity profile A_i_(x) = A(x, t_i_) was generated. The profiles A_i_(*x*) were normalized to the maximal value of the first profile A_1_ = max {A_1_(*x*)} and smoothed using the exponential moving average with the smoothing constant 0.2. The sequence of the normalized smoothed profiles A^s^_i_(*x*) = A_i_(*x*)/A_1_ corresponds to a surface which was presented as a heat map showing the levels A^s^ = const. The length of each profile di corresponds to the ring perimeter and the time ordered sequence of d_i_ produces the perimeter dynamics.

## Acknowledgments

We thank Francisco Calheiros for fruitful discussions and critical reading of the manuscript. This project has received funding from the European Research Council under the European Union’s FP7 ideas programme (grants 260892 and 338410) and Horizon 2020 research and innovation programme (grant 640553), FEDER funds through the Operational Competitiveness Programme-COMPETE, national funds through FCT-Funda9&o para a Ciencia e a Tecnologia under the project FC0MP-01–0124-FEDER-028255 (PTDC/BEX-BCM/0654/2012), FLAD Life Science 2020 and The Louis-Jeantet Young Investigator Award (to HM). AXC, RG and IAT have FCT Investigator positions funded by national funds through FCT and co-funded by the European Social Fund (ESF) through the Human Potential Operational Programme (POPH) Type 4.2-Promotion of Scientific Employment (IF/00901/2013/CP1157/CT0001, IF/01015/2013/CP1157/CT0006 and IF/00082/2013, respectively). AMS holds a postdoctoral fellowship by FCT (SFRH/BPD/95707/2013). DO received funding from the Project “NORTE-07–0124-FEDER-000003-Cell Homeostasis Tissue Organization and Organism Biology” cofunded by Programa Operacional Regional do Norte (ON.2—O Novo Norte), under the Quadro de Referenda Estrategico Nacional (QREN), through the Fundo Europeu de Desenvolvimento Regional (FEDER) and by FCT.

## List of abbreviations

CI: confidence interval

## Supplemental Figures and Supplemental Figure Legends

**Figure S1.**
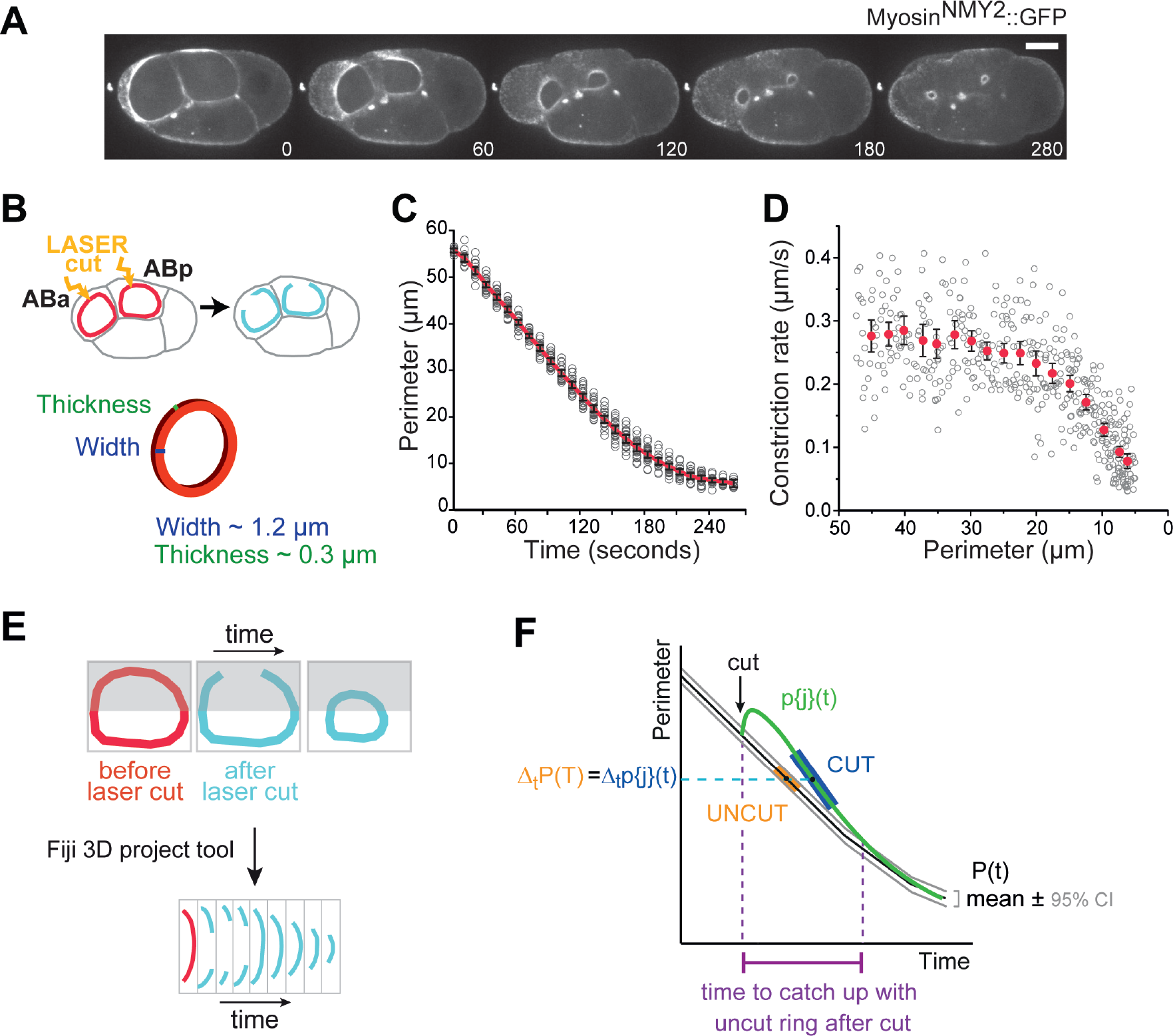
End-on view time-lapse imaging of the contractile ring of AB cells in the 4-cell *C. elegans* embryo. **A.** End-on view of constricting contractile rings in a 4-cell *C. elegans* embryo expressing myosin^NMY–2^::GFP. The ABa and ABp cells undergo ring constriction parallel to the imaging plane. Time is in seconds. Scale bar, 10 μm. **B.** Schematic of 4-cell embryo illustrating where laser cuts were inflicted. The contractile ring in the early *C. elegans* embryo is estimated to be 1.3 μm wide and 0.3 μm thick (Carvalho et al., 2009). **C.** Perimeter versus time plot for contractile ring constriction in ABa and ABp cells (n=22). Red line represents the average perimeter with black error bars denoting the 95% confidence interval (CI) of the mean. Gray circles show individual values. **D.** Mean constriction rate (red dots with error bars representing 95% CI; see methods for details) plotted against decreasing ring perimeter in ABa and ABp cells (n=22). Grey circles show individual values. **E.** Schematic illustrating how kymographs of the cut region were generated for Fig. 1B–D and Fig. 2A. **F.** Schematic illustrating how instantaneous rates of perimeter change in figure 3B were calculated.

**Figure S2.**
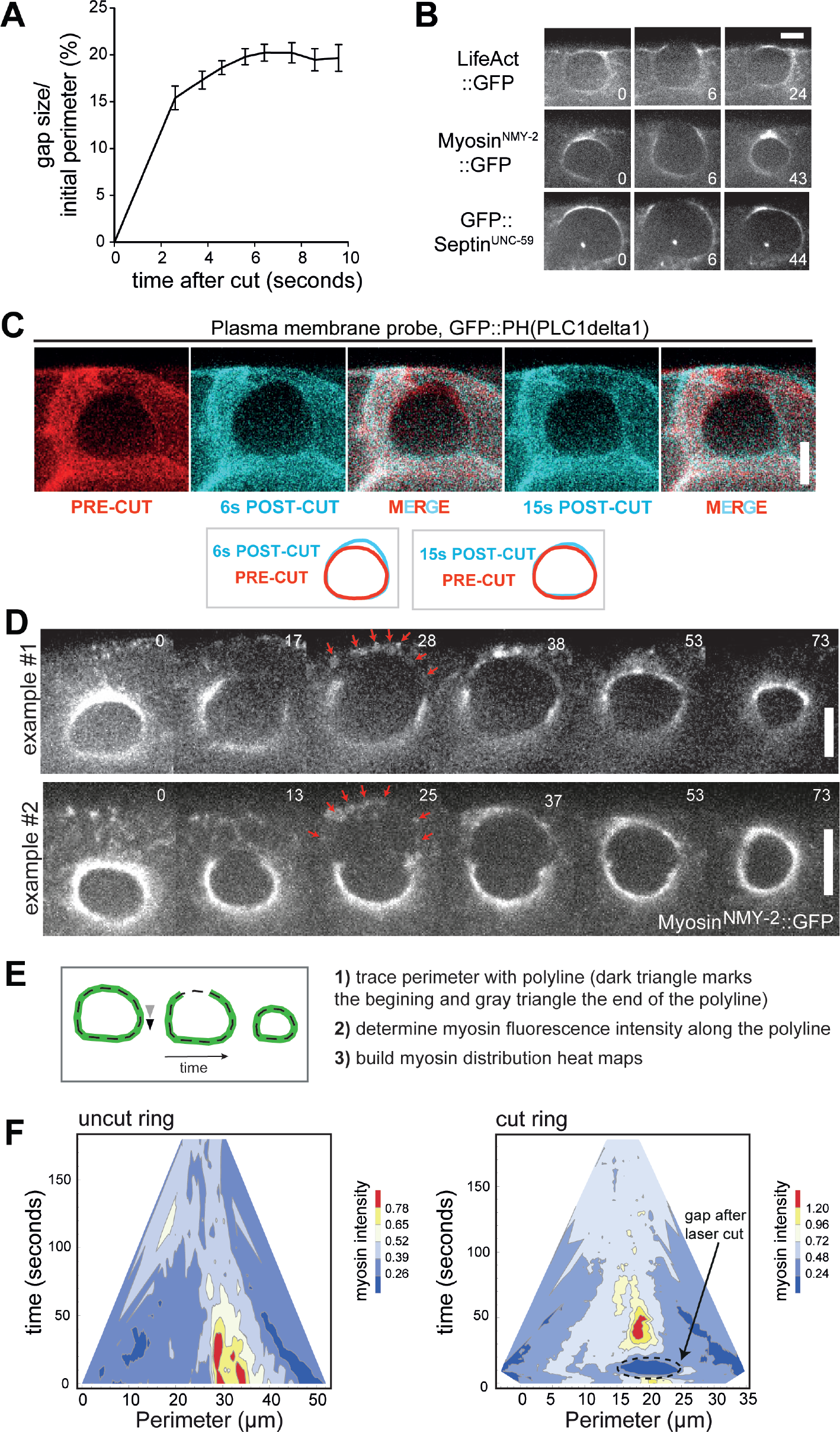
Characterization of gap formation and gap repair in embryos expressing fluorescent probes for the contractile ring and plasma membrane. **A.** Gap size normalized to initial perimeter during the first 10 s after the laser cut. Data for all examples presented in figure 2C is shown (n=24). **B.** Time lapse stills of laser microsurgery experiments performed in embryos expressing fluorescent probes for actin (lifeAct::GFP), nonmuscle myosin II (NMY-2::GFP), and one of the two septins (UNC-59::GFP). **C.** Stills of a constricting ring cut by laser microsurgery in an embryo expressing a pleckstrin homology domain (PH) fused to GFP to mark the plasma membrane. Overlay at the bottom compares the ring outlines before the cut against the ring outline 6 s post-cut (local deformation of plasma membrane) and 15 s post-cut (plasma membrane deformation no longer evident). D. Stills of two cut rings that fail the first attempt of repair. Red arrows point to foci of myosin^NMY-2^::GFP that accumulates de novo along the length of the gap. Time is in seconds and time zero corresponds to the frame before the laser cut. E. Schematic illustrating how heat maps were generated that show the myosin^NMY-2^::GFP distribution along the contractile ring circumference over time. F. Heat maps for myosin^NMY-2^::GFP as described in (E). Scale bars, 5 μm.

**Figure S3.**
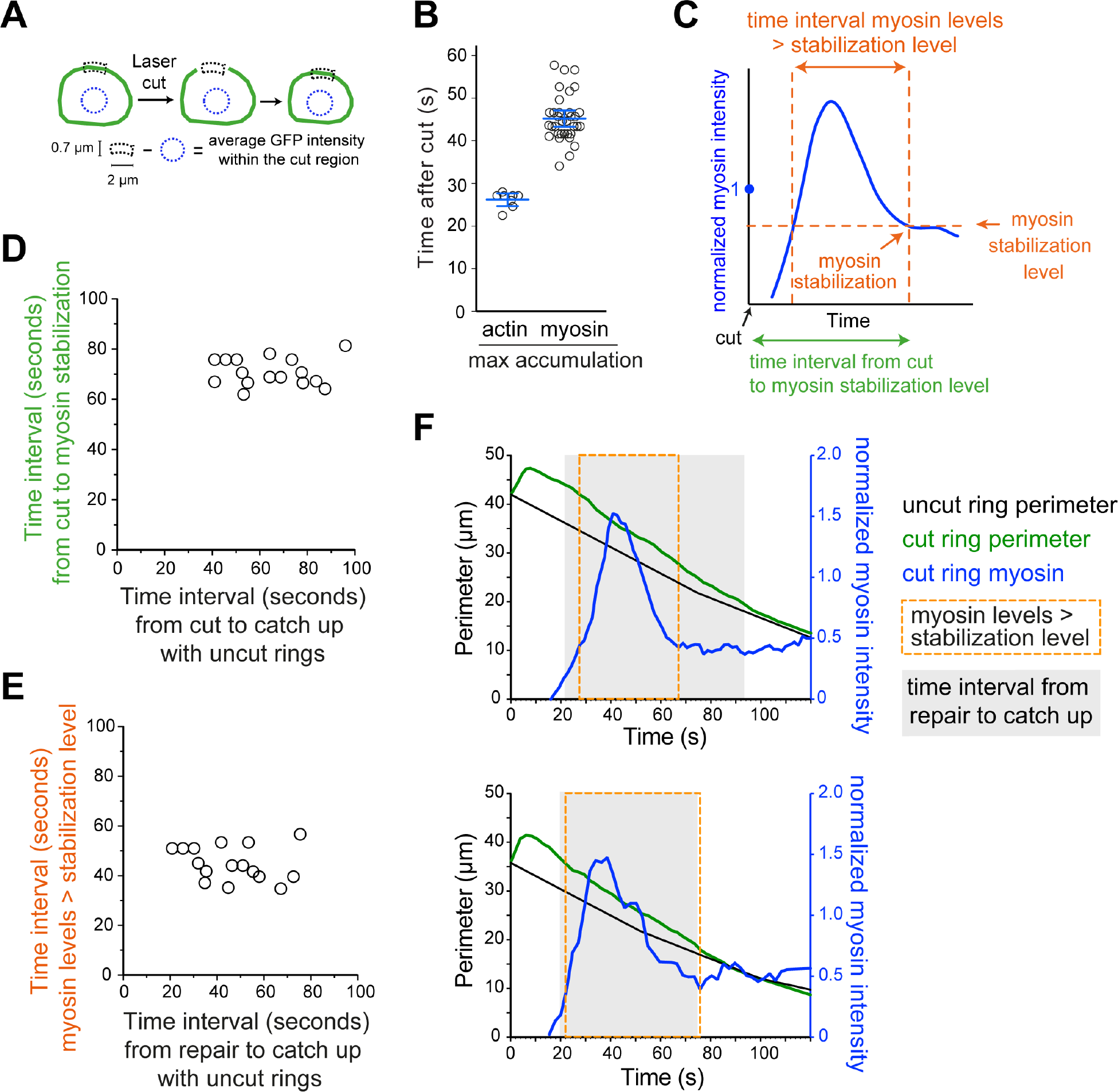
Characterization of myosin hyper-accumulation after gap repair A. Schematic illustrating the quantification method for average myosin intensity levels in the cut region. A region of constant dimensions was placed over the severing region at time points just before and after the cut. Cytoplasmic signal was subtracted and values were normalized to the average myosin intensity before the cut. **B.** Time period between laser cut and maximum accumulation of actin (lifeAct::GFP) or myosin (NMY2::GFP) in the repaired region. Error bars represent the 95% CI of the mean. **C.** Schematic illustrating how time intervals plotted in D, E, and F were determined. **D.** Time interval between laser cut and stabilization of myosin levels versus time until cut rings caught up with perimeters of uncut rings (n=16, Pearson correlation test not significant). **E.** Time interval during which myosin levels were above the stabilization level versus time interval between completion of repair and catching up (n=16, Pearson correlation test not significant). **F.** Two representative examples of perimeter (green) and normalized myosin levels (blue) versus time in cut rings. Average perimeter of uncut rings is shown in black. Normalized myosin levels in uncut rings, which remain constant throughout constriction, are not shown. The grey area marks the time interval between completion of repair and catching up. The orange box marks the time during which myosin levels are above the stabilization level, as depicted in C. Top graph shows an example of a cut ring in which myosin levels stabilize before the ring perimeter catches up with that of uncut rings. The bottom graph shows an example in which myosin levels drop coincident with ring perimeter catching up with that of uncut rings.

## Table and Table Legend

**Table S1.** List of the worm strains used in this study.

| Strains | Genotype |
| --- | --- |
| N2 | Ancestral |
| OD822 | unc-119(ed3)III; ltSi201[pAC65; Pnmy-2::GFP; cb-unc-119(+)]II |
| GCP21 | ltIs157 [pAC16;pie-1/Life-Act::GFP; unc-119 (+)]; ltIs37 [pAA64; pie-1/mCherry::his-58; unc-119 (+)] IV |
| GCP57 | unc-119(ed3)III; prtSi2[pAC71; Pnmy-2:nmy-2 re-encoded::mCherry::StrepTagII::3'UTRnmy-2; cb-unc-119(+)]II + unc-119(ed3) III; ltIs44 [pAA173; pie-1/GFP::PH(PLC1delta1); unc-119 (+)] |
| OD121 | unc-119(ed3) III; ltIs20[pASM10; pie-1/GFP::unc-59; unc-119(+)]; ltIs37 [pAA64; pie-1/mCherry::his-58; unc-119 (+)] IV |

## References

Audhya,A. 2005. A complex containing the Sm protein CAR-1 and the RNA helicase CGH-1 is required for embryonic cytokinesis in Caenorhabditis elegans. J Cell Biol. 171:267–279. doi:10.1083/jcb.200506124.

Bement,W.M., and D.G. Capco. 1991. Analysis of inducible contractile rings suggests a role for protein kinase C in embryonic cytokinesis and wound healing. Cell Motil Cytoskeleton. 20:145–157. doi:10.1002/cm.970200207.

Biro,M., Y. Romeo, S. Kroschwald, M. Bovellan, A. Boden, J. Tcherkezian, P.P. Roux, G. Charras, and E.K. Paluch. 2013. Cell cortex composition and homeostasis resolved by integrating proteomics and quantitative imaging. Cytoskeleton. 70:741–754. doi:10.1002/cm.21142.

Carvalho,A., A. Desai, and K. Oegema. 2009. Structural memory in the contractile ring makes the duration of cytokinesis independent of cell size. Cell. 137:926–937. doi:10.1016/j.cell.2009.03.021.

Carvalho,A., S.K. Olson, E. Gutierrez, K. Zhang, L.B. Noble, E. Zanin, A. Desai, A. Groisman, and K. Oegema. 2011. Acute Drug Treatment in the Early C. elegans Embryo. PLoS ONE. 6:e24656. doi:10.1371/journal.pone.0024656.

Collinet,C., M. Rauzi, P.-F. Lenne, and T. Lecuit. 2015. Local and tissue-scale forces drive oriented junction growth during tissue extension. Nat Cell Biol. 17:1247–1258. doi:10.1038/ncb3226.

Colombelli,J., A. Besser, H. Kress, E.G. Reynaud, P. Girard, E. Caussinus, U. Haselmann, J.V. Small, U.S. Schwarz, and E.H.K. Stelzer. 2009. Mechanosensing in actin stress fibers revealed by a close correlation between force and protein localization. J Cell Sci. 122:19281928. doi:10.1242/jcs.054577.

Founounou,N., N. Loyer, and R. Le Borgne. 2013. Septins Regulate the Contractility of the Actomyosin Ring to Enable Adherens Junction Remodeling during Cytokinesis of Epithelial Cells. Developmental Cell. 24:242–255. doi:10.1016/j.devcel.2013.01.008.

Green,R.A., E. Paluch, and K. Oegema. 2011. Cytokinesis in Animal Cells. Annu Rev Cell Dev Biol. 28:120717164503001. doi:10.1146/annurev-cellbio-101011-155718.

Guillot,C., and T. Lecuit. 2013. Adhesion Disengagement Uncouples Intrinsic and Extrinsic Forcesto Drive Cytokinesis in Epithelial Tissues. Dev Cell. 24:227–241. doi:10.1016/j.devcel.2013.01.010.

Herszterg,S., A. Leibfried, F. Bosveld, C. Martin, and Y. Bellaiche. 2013. Interplay between the Dividing Cell and Its Neighbors Regulates Adherens Junction Formation during Cytokinesis in Epithelial Tissue. Developmental Cell. 24:256–270. doi:10.1016/j.devcel.2012.11.019.

Kamasaki,T., M. Osumi, and MabuchiII. 2007. Three-dimensional arrangement of F-actin in the contractile ring of fission yeast. J Cell Biol. 178:765–771. doi:10.1083/jcb.200612018.

Kumar,S., I.Z. Maxwell, A. Heisterkamp, T.R. Polte, T.P. Lele, M. Salanga, E. Mazur, and D.E. Ingber. 2006. Viscoelastic Retraction of Single Living Stress Fibers and Its Impact on Cell Shape, Cytoskeletal Organization, and Extracellular Matrix Mechanics. Biophys J. 90:37623773. doi:10.1529/biophysj.105.071506.

Mandato,C.A., and W.M. Bement. 2001. Contraction and polymerization cooperate to assemble and close actomyosin rings around Xenopus oocyte wounds. J Cell Biol. 154:785–797. doi:10.1083/jcb.200103105.

Munjal,A., J.-M. Philippe, E. Munro, and T. Lecuit. 2015. A self-organized biomechanical network drives shape changes during tissue morphogenesis. Nature. 524:351–355. doi:10.1038/nature 14603.

Sain,A., M.M. Inamdar, and F. Julicher. 2015. Dynamic Force Balances and Cell Shape Changes during Cytokinesis. Phys. Rev. Lett. 114:048102.

Schindelin,J., I. Arganda-Carreras, E. Frise, V. Kaynig, M. Longair, T. Pietzsch, S. Preibisch, C. Rueden, S. Saalfeld, B. Schmid, J.-Y. Tinevez, D.J. White, V. Hartenstein, K. Eliceiri, P. Tomancak, and A. Cardona. 2012. Fiji: an open-source platform for biological-image analysis. NatMeth. 9:676–682. doi:10.1038/nmeth.2019.

Schneider,C.A., W.S. Rasband, and K.W. Eliceiri. 2012. NIH Image to ImageJ: 25 years of image analysis. Nat Meth. 9:671–675.

Stachowiak,M.R., C. Laplante, H.F. Chin, B. Guirao, E. Karatekin, T.D. Pollard, and B. O'Shaughnessy. 2014. Mechanism of Cytokinetic Contractile Ring Constriction in Fission Yeast. Developmental Cell. 29:547–561. doi:10.1016/j.devcel.2014.04.021.

Tinevez,J.-Y., U. Schulze, G. Salbreux, J. Roensch, J.-F. Joanny, and E. Paluch. 2009. Role of cortical tension in bleb growth. Proceedings of the National Academy of Sciences. 106:18581–18586. doi:10.1073/pnas.0903353106.

Turlier,H., B. Audoly, J. Prost, and J.-F. Joanny. 2014. Furrow Constriction in Animal Cell Cytokinesis. Biophys J. 106:114–123. doi:10.1016/j.bpj.2013.11.014.

Zhang,W., and D.N. Robinson. 2005. Balance of actively generated contractile and resistive forces controls cytokinesis dynamics. Proc Natl Acad Sci USA. 102:7186–7191. doi:10.1073/pnas.0502545102.

